# COFFEE-PRESC: A fast pre-screening method using compound retrieval by pairwise positional relationship of representative fragments

**DOI:** 10.64898/2025.12.07.692874

**Authors:** Masayoshi Shimizu, Satoshi Yoneyama, Keisuke Yanagisawa, Yutaka Akiyama

## Abstract

Protein-ligand docking is one of the most widely used methods in structure-based virtual screening in the early stages of drug discovery. Its calculations require approximately one minute per compound, making exhaustive evaluation of ultra-large libraries containing billions of molecules computationally impractical.

In this study, we propose COFFEE-PRESC (COmpound Filtering by Fragment pair-based Efficient Evaluation for PRE-SCreening), a fast, fragment-based pre-screening method. COFFEE-PRESC first docks fragments in a pre-constructed fragment set to the target protein and enumerates multiple favorable protein-fragment docking poses and then pairs them to consider pairwise positional relationship. The fragment set is composed of a small number of representative fragments that exhibit high similarity to many other fragments, enabling coverage of a large and diverse chemical space.

Compounds that contain stuructures similar to fragment pairs are then retrieved through similarity-based searches. This retrieval methodology guarantees that the mutual positional relationship of the two matched fragments does not spatially collide.

Finally, the retrieved compounds are evaluated using docking scores of the representative fragments and similarity values between the representative and individual fragments matched in the compound retrieval process. COFFEE-PRESC was 32-fold faster while higher accuracy than Spresso, a existing pre-screening tool, highlighting its potential for application to ultra-large compound library screening.

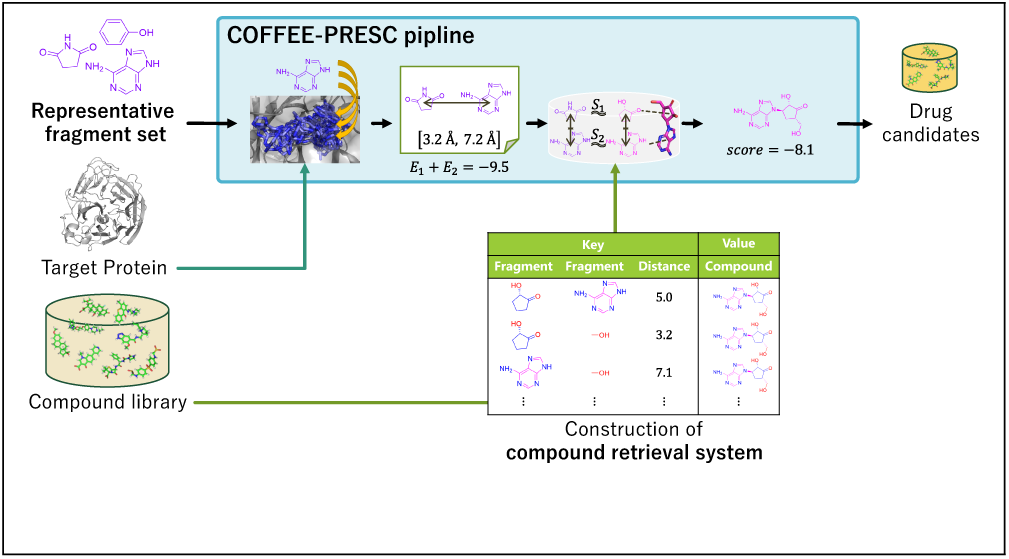
TOC Graphic.

## Introduction

Structure-based virtual screening (SBVS), which selects candidate compounds using the tertiary structures of target proteins, is widely employed in the early stages of drug discovery. In recent years, the size of compound libraries used in SBVS has increased exponentially; for example, ZINC22 contains more than 37 billion compounds. ^1^ In such cases, proteinligand docking (hereafter referred to as docking), which sequentially predicts the structure of protein-ligand complexes, becomes computationally infeasible for evaluating all compounds. For instance, AutoDock Vina^2,3^ requires approximately one minute per compound; therefore, screening the entire ZINC22 database takes approximately 70,000 CPU core years. To address this issue, compound libraries are typically filtered before docking through prescreening methods based on physico-chemical features, such as Lipinski’s rule of five^4^ and the quantitative estimate of drug-likeness (QED).^5^ This strategy enables rapid filtering of compounds;^6^ however, simply applying it will not filter through the vast amount of data, and if we drastically reduce the number of compounds, compound diversity will decrease unacceptably. This would compromise one of the main advantages of SBVS, which is its ability to discover novel chemotypes. Meanwhile, active learning methods that iteratively

select compounds to dock based on surrogate model predictions and uncertainties have been proposed.^7^ However, this strategy is also expected to reduce chemical diversity when it relies too heavily on the assumptions of the surrogate model. To maintain chemical diversity, it is crucial to thoroughly evaluate all compounds using structure-based methods. To overcome the limitation, several methods have been proposed that incorporate protein structural information to accelerate compound evaluation, including simplified docking approaches.^8–10^ Nevertheless, faster methods are still required for such billions class compound libraries.

Another acceleration strategy, fragment-based virtual screening (FBVS), has been introduced. FBVS decomposes chemical compounds into fragments (smaller substructures). Compounds are evaluated based on the docking results of their constituent fragments. Since the number of unique fragments is far smaller than the number of compounds, FBVS can significantly reduce the number of required docking calculations. Yanagisawa *et al.*^9^ reported that 28,629,602 compounds from the ZINC12 database could be decomposed into only 263,319 unique fragments; thus, fragment decomposition is theoretically 100-fold faster or more. Several fragment-based methods have been developed. For example, FlexX ^11^ and DOCK 6^12,13^ reconstruct compounds incrementally by extending from a core fragment docking pose with the target protein. eHiTS^14^ docks a compound by combinatorial optimization, which is formulated as a problem to select one possible fragment docking pose for each fragment using a maximum clique search on a graph whose nodes correspond to fragment docking poses. REstretto^15^ uses pre-generated compound conformers and evaluates fragments with simplified scoring. However, these methods are computationally expensive due to the approach that estimates the binding modes of each compound while accounting for their numerous conformations. To maximize the advantage of the reduced number of fragments, it is crucial to minimize the computational cost that scales with the number of compounds; thus, a fragment-based pre-screening method, Spresso,^9^ was proposed. Spresso significantly reduces the computational cost of the compound evaluation step by simply summing up fragment-docking scores without considering the mutual positional relationship between fragments. Spresso is approximately 200-fold faster than the commercial software, Glide (HTVS mode),^16^ on approximately 29 million compounds. However, because only the best docking score for each fragment is considered, the fragment docking poses are ignored. As a result, Spresso entirely ignores mutual positional relationships between the best-scoring fragment poses. This can lead to implausible combinations in which good-scoring fragment docking poses collide, resulting in inappropriate evaluation for compounds that contain multiple core fragments.

In this study, we propose COFFEE-PRESC (COmpound Filtering by Fragment pair-based Efficient Evaluation for PRE-SCreening), a novel and efficient fragment-based prescreening method. COFFEE-PRESC retrieves compounds using a pre-constructed database through a lookup based on the favorable pairwise positional relationship of fragments composing the compounds. Considering the mutual positional relationships between fragments improves screening accuracy. Furthermore, to enable ultra-large virtual screening, it evaluates diverse fragments based on their similarity while reducing computational costs by using only the structures of a small number of pre-selected representative fragments for docking. Our method offers fast pre-screening that considers pairwise positional relationships between fragments and is expected to be applicable to ultra-large compound libraries.

## Materials and methods

Considering the mutual positional relationships between fragments should be essential to improve pre-screening accuracy compared with Spresso, while maintaining computational efficiency is also crucial. To this end, we introduced two strategies. First, to consider the relative positions of fragments in rapid compound evaluation, we adopted a database lookup of compounds using mutual positional relationship of two fragments. Our approach retrieves compounds based on the set of fragments and their all-pair distances. It skips explicit compound reconstruction while considering all pairwise positional relationships of fragments. Although information retrieval is fast, this approach is slightly slower than the fragment docking score summation of Spresso. Second, we leveraged fragment similarity to reduce computational cost. Molecular structural similarity correlates with the similarity of bioactivity^17^ and docking scores. In addition, recent studies have demonstrated that similarity-based screening is effective at the fragment level. ^18^ This suggests that defining a set of representative fragments that capture similarity to many other fragments can reduce fragment docking, resulting in reduced computational cost.

The aforementioned two strategies constitute our proposed pre-screening method, COFFEEPRESC. COFFEE-PRESC first perform docking calculations using a small number of representative fragments as input. It then enumerate promising fragment pairs from fragment docking results and perform high-throughput compound retrieval, considering fragment similarity.

Figure 1 shows the workflow of COFFEE-PRESC along with its key strategies. We describe the construction of the representative fragment set and compound retrieval system and then present the overall COFFEE-PRESC pipeline.

**Figure 1:**
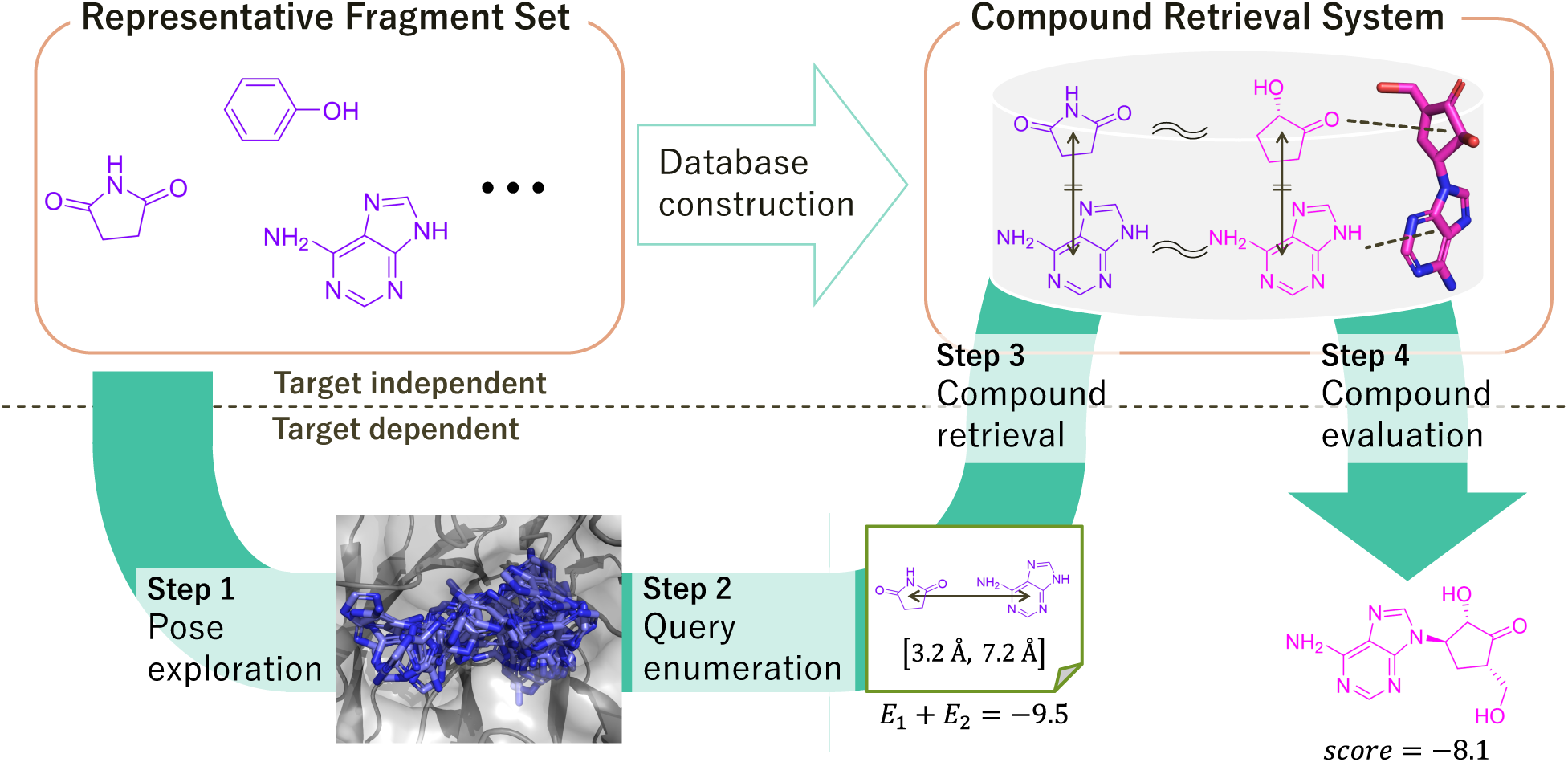
COFFEE-PRESC workflow. The method consists of four steps: pose exploration, query enumeration, compound retrieval, and compound evaluation.

### Representative fragment set

COFFEE-PRESC requires a representative fragment set as input, in addition to the target protein and compound library to search. We show an example of creating such a fragment set. The fragments contained in the representative fragment set should be similar to as many outside fragments as possible. Selecting fragments that are similar to many others enables the use of a smaller fragment set.

We first constructed an initial fragment population and then selected representatives from this population. As the compound population for selecting representative fragments, we used the DrugBank Approved subset^19^ (Release 5.1.10, 2023-01-04), which contains 2,587 FDA-approved compounds. To use the scoring function of AutoDock Vina, we retained compounds composed only of H, C, N, P, O, S, F, Cl, Br, and I, without isotopes, and containing at least one carbon atom. This filter removed 324 compounds, leaving 2,263 compounds. We then decomposed these compounds using the Spresso protocol,^9^ resulting in 1,620 unique fragments used for representative fragment selection.

From the generated fragment population, we selected 50 fragments as a representative fragment set. Assuming that fragments are relatively small, rigid substructures, we first filtered out fragments that are unsuitable as representatives. We excluded fragments that met any of the following:

1. contain any ring with ≥ 7 members,
2. contain ≥ 3 non-aromatic rings,
3. contain ≥ 4 rings,
4. contain ≥ 7 non-ring heavy atoms, or
5. consist of only one heavy atom.

This removed 394 fragments, yielding 1,226 fragments. We then clustered the filtered fragments using Ward’s method. For the clustering, we used the distance matrix *D_ij_* defined from a fragment similarity value as

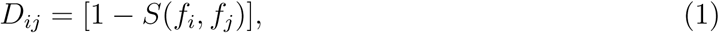

where *f_i_* and *f_j_* are fragments of interest, and *S*(*f_i_, f_j_*) ∈ [0, 1] denotes the fragment similarity value. We adopted the similarity measure proposed by Yoshimori *et al.*^20^ The number of clusters corresponded to the number of representative fragments. From each cluster *C*, we selected one representative fragment as follows. For each fragment *f_i_* ∈ *C*, we computed the mean intra-cluster distance, 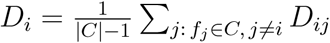 The fragment with the smallest mean distance was chosen as the representative of that cluster.

### Compound retrieval system

We describe the compound retrieval system based on fragment pairs―another key strategy of COFFEE-PRESC. Each fragment composing a compound possesses spatial constraints to avoid collisions. It is possible to improve pre-screening accuracy by considering mutual positional relationships between fragments; these were not considered in Spresso when calculating compound scores. Therefore, we propose a fragment pair-based compound retrieval system that returns compounds consistent with the mutual positional relationship of two fragments given in a query. The compound retrieval system comprises two stages: database construction and compound retrieval stages.

### Database construction

The database construction stage can be divided into four steps (Figure 2a): fragment decomposition, generating conformers of compounds, fragment pair enumeration, and database registration. First, compounds are decomposed using the Spresso protocol.^9^ Thereafter, three-dimensional compound conformers are generated by a conformer generator, such as OMEGA,^21^ to calculate the three-dimensional distance between fragments constituting the compound.

**Figure 2:**
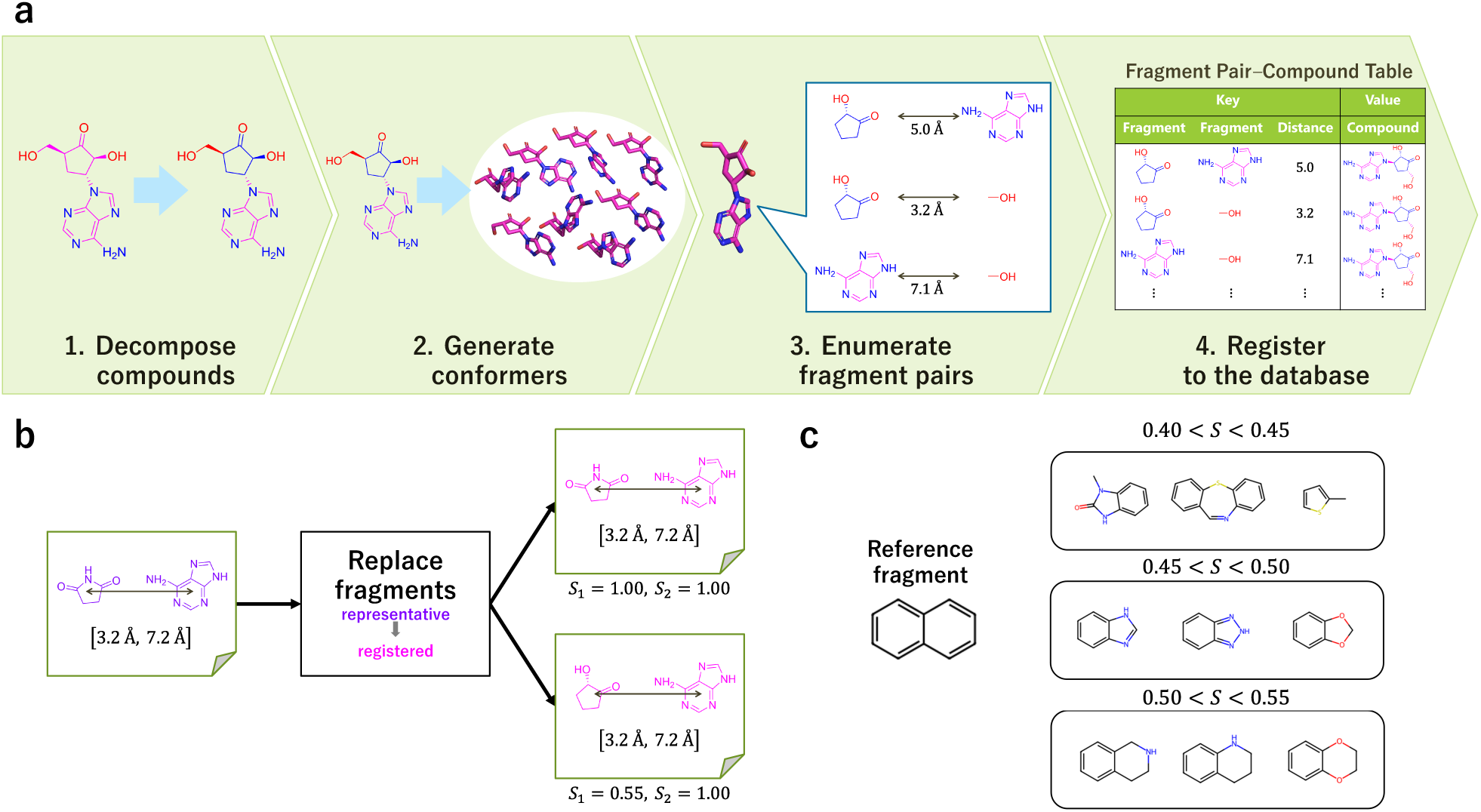
(a) Database construction for the compound retrieval system. (b) Replacing each representative fragment in the query with similar registered fragments. (c) Specific examples of the relationship between fragment structures and similarity values.

After conformer generation, the fragment pair-compound table is created based on the fragment information associated with the conformers. Note that the conformers are registered at this time as compound information represented in formats like SMILES. The fragment pair-compound table has a key–value structure. The key consists of the two fragments and the inter-centroid distance between them, while the value is the corresponding compound. The fragments are referred to as registered fragments. Note that identical keys may be found multiple times in the table. This is because within multiple conformers of a compound, or even in different compounds, structures may exist where the distance between two fragments is similar. To register a compound to the table, all possible fragment pairs are enumerated and registered. If a compound contains *n* ≥ 2 fragments, *_n_*C_2_ keys are registered. For storage efficiency, each of the two fragments is assigned integer IDs, and inter-centroid distances are quantized in 0.1 Å increments. The key is encoded in a 64-bit unsigned integer: 27 bits for the fragment ID and 10 bits for the quantized distance (total 27 × 2 + 10 = 64 bits). After registering all compounds, the table is sorted in ascending order based on the key. Since the distance is stored in the last 10 bits of the 64-bit key, the keys registered in the table are grouped by fragment type and sorted by distance.

### Compound retrieval

In compound retrieval, we use a single fragment pair query as the input, which consists of two representative fragments and the distance between them. At this stage, by considering similarity, compounds with similar fragments can be retrieved. Furthermore, since fragment pair queries cannot be enumerated exhaustively and exact compound matching based on strict inter-fragment distances is impractical, we also enable retrieving compounds with a certain margin in inter-fragment distances. Therefore, a query is defined as a pair of representative fragments together with an allowable range of inter-centroid distances (minimum and maximum). As in database construction, distances are quantized at 0.1 Å. The queries are also enumerated considering fragment similarity; as shown in Figure 2b if the first representative fragment of a query has two similar registered fragments, the query is bifurcated to two queries. We recommend treating two fragments as similar fragments if their similarity value *S*(*f_i_, f_j_*) is above 0.45 using the Yoshimori’s similarity measure,^20^ from the standpoint of both retrieval speed and diversity of matched fragments. Several examples of similarity values are shown in Figure 2c. Compounds are then retrieved using enumerated queries. Within the table, records for each fragment type are sorted by distance. Therefore, performing two binary search sorts per query allows us to enumerate compounds that satisfy these conditions. For each matched compound, the system outputs two fragment similarity values (*S*_1_*, S*_2_) corresponding to the two representative fragments in the query and their matched registered fragments.

### COFFEE-PRESC pipeline

We have described the two key strategies of COFFEE-PRESC, namely the representative fragment set and the compound retrieval system. We introduce the workflow of COFFEEPRESC, our pre-screening method (Figure 1). The input consists of a target protein, a cuboid region of a binding site on it, a compound library to screen, and representative fragment set. The output is a ranked list of candidate compounds retrieved by the compound search, together with pre-screening scores, which are then used to select compounds for subsequent time-consuming but more accurate screening steps.

### Step 1: Docking pose exploration of representative fragments

First, COFFEE-PRESC enumerates candidate fragment docking poses for each representative fragment. Each fragment is placed around the docking region, at equal intervals (0.25 Å increments as a default) translation and dodecahedron-based rotation, and each docking pose is evaluated using the AutoDock Vina energy score function.^2^ Candidate fragment docking poses are greedily selected from high-scoring fragment docking poses, and the centroids of these poses and their scores are stored. The docking score is denoted by *E*. Note that these docking poses are used for similarity-based compound retrieval in Step 3; however, each docking pose is a good pose for its representative fragment and not necessarily for similar fragments. Therefore, the docking scores of similar but different fragments are likely to be worse. The number of candidate fragment docking poses (*N*) per representative fragment is parameterized. To select various fragment docking poses, a minimum distance (*r_e_*) between candidate poses is also parameterized.

### Step 2: Query enumeration

Second, fragment pair queries are enumerated with all the candidate fragment docking poses. For example, with 50 representative fragments and *N* = 10 candidates per fragment (i.e., 500 candidates in total), _500_C_2_ queries are generated. Each query consists of two representative fragments and their inter-centroid distance. Each query has the sum of the docking scores of the corresponding candidate poses, *E*_1_ + *E*_2_. Because each candidate is selected with a minimum distance *r_e_*, the inter-centroid distance of the query should have an interval that reflects that. If the observed distance is *d* Å, the interval is set to [*d* − 2*r_e_, d* + 2*r_e_*].

Different queries with the same two fragments can have an intersection of inter-centroid distance interval. For example, the intervals [2.1 Å, 6.1 Å] and [3.2 Å, 7.2 Å] have an intersection of [3.2 Å, 6.1 Å]. If this intersection is ignored, compounds will be retrieved multiple times under the same conditions, resulting in redundant computational costs. To eliminate redundancy before retrieval, we keep the best (lowest) *E*_1_ + *E*_2_ for each duplicated condition, as illustrated in Figure 3. Hereinafter, this fragment pair query after unification is simply referred to as a query.

**Figure 3:**
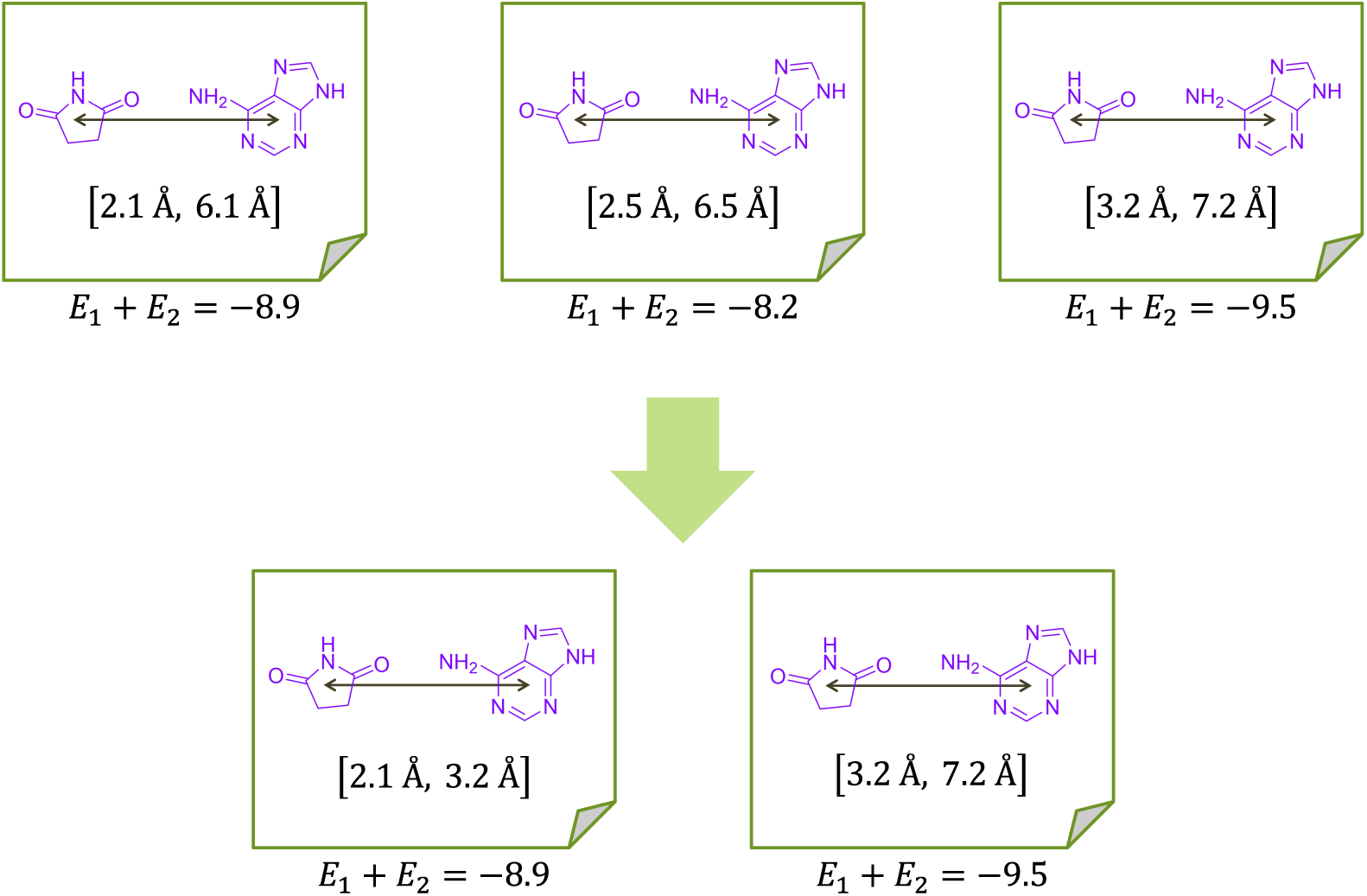
Unification of duplicate fragment pair queries (keeping the lowest sum of docking scores *E*_1_ + *E*_2_).

### Step 3: Compound retrieval

COFFEE-PRESC then retrieves the compounds with each query, obtaining matched compounds together with their similarity values (*S*_1_ and *S*_2_).

### Step 4: Compound evaluation

Assuming that only certain fragments within a compound dominate binding affinity in many cases, the compound score can be roughly estimated using only the two fragment docking scores in the query. Thus, the sum of fragment docking scores *E*_1_ +*E*_2_ is used as a compound score. However, the matched registered fragments can differ from the query representative ones, and fragment docking scores can also differ. As described in Step 1, when the matched registered fragments are superimposed onto the docking poses of the query representative fragments, the former are likely to yield poorer docking scores than the latter. Moreover, a lower similarity value is likely to result in a larger discrepancy between the docking scores. Thus, to reflect this tendency, the score of compound fragment *E*^′^ should be penalized from the docking score of the query fragment *E* according to similarity value *S*:

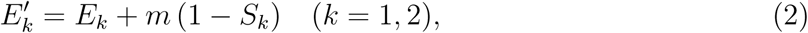

where *m* ≥ 0 is a penalty coefficient. A linear penalty was chosen because preliminary experiments suggested a somewhat linear relationship between the similarity value *S* and the difference in docking scores *E*^′^ − *E* for two fragment with high similarity. Therefore, the compound score can be expressed as follows:

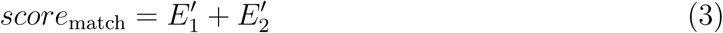

In Step 3, compounds matched to each query are output individually, often resulting in the same compound being listed multiple times. In such cases, only the best *score*_match_ is retained so that each compound is associated with a single compound score.

### Dataset

We used the Directory of Useful Decoys, Enhanced (DUD-E) ^22^ to evaluate the pre-screening performance during virtual screening. DUD-E provides 102 targets, each comprising a cocrystal structure, active compounds, and decoy compounds. On average, each target contains 14,059 compounds, with an active:decoy ratio of roughly 1:50. Compounds for each target were preprocessed using LigPrep (Schrödinger Suite, version 2024-1), and conformers of compounds were generated using OMEGA^21^ (version 5.0.0.3), both with default parameters. We constructed 50 representative fragments using the aforementioned method.

### Metrics

We assessed both screening accuracy and execution time. Accuracy was measured by the enrichment factor (EF):

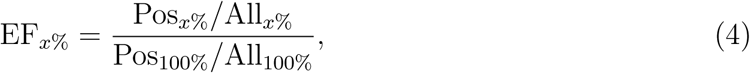

where Pos*_x_*_%_, All*_x_*_%_, Pos_100%_, and All_100%_ represent the number of active compounds in the top *x*% of screened compounds, the number of compounds in the top *x*% of screened compounds, the total number of active compounds, and the total number of compounds, respectively. Note that EF*_x_*_%_ = 1.0 means a random prediction. We calculated EF_1%_ and EF_2%_. Execution time was measured in terms of CPU core time (recorded with the Linux time command).

### Performance comparison with other methods

Pre-screening methods such as COFFEE-PRESC are not intended to complete virtual screening on their own. Therefore, a practical evaluation must involve not only pre-screening methods but also the following compound docking calculation. As baselines for pre-screening, we used Spresso and random sampling. Compound docking was performed using AutoDock Vina. The three-step procedure used to evaluate accuracy was as follows: (i) with each pre-screening method, 2%, 5%, or 10% of the number of all target compounds were selected; (ii) pre-screened candidates were docked using AutoDock Vina^2,3^ (version 1.2.7) to obtain the docking score; and (iii) EF_1%_ and EF_2%_ were calculated based on the top 1% and 2% of compounds ranked by the docking scores.

### Experimental setups

The center coordinates and the lengths on each side of the docking region boxes were determined using eBoxSize.^23^ The COFFEE-PRESC parameters selected in the experiments are shown in Table 1, which were determined from preliminary experiments on 12 DUD-E targets listed in Table S1. The number of candidate fragment docking poses per representative fragment, *N* , was set to 40: beyond this value, screening accuracy saturated while computational cost continued to grow. The minimum distance between candidate fragment docking poses *r_e_* was set to 1.0 Å, following the default minimum RMSD difference between docking poses in AutoDock Vina. The compound retrieval database was constructed independently for each target, registering compounds of the actives and decoys for the target. For compound docking using AutoDock Vina, we used default settings with the seed fixed at 42. For Spresso, we reimplemented the compound evaluation step and the compound scoring function followed GS_3_.^9^

**Table 1:** COFFEE-PRESC Parameters

| Parameter | Description | Value |
| --- | --- | --- |
| $N$ | Number of candidate docking poses per representative fragment | 40 |
| $m$ | Similarity-based penalty coefficient in compound scoring function | 8 |

### Computing environment

All calculations were conducted on the TSUBAME4.0 supercomputing system at Institute of Science Tokyo, Japan. Each node comprised two AMD EPYC 9654 CPUs (96 cores per CPU) and 768 GB RAM. AutoDock Vina was executed using 8 CPU cores; other tools were run with a single CPU core.

## Results

### Representative fragment set

The 50 representative fragments constructed in this study are shown in Figure 4. Among them, five contain no rings; the numbers of fragments containing one, two, and three rings are 28, 14, and 3, respectively. In addition, 29 fragments contain at least one aromatic ring. Fragments containing oxygen, nitrogen, and halogens are found 27, 30, and 7 times, respectively, suggesting that substructures commonly involved in protein interactions are well represented.

**Figure 4:**
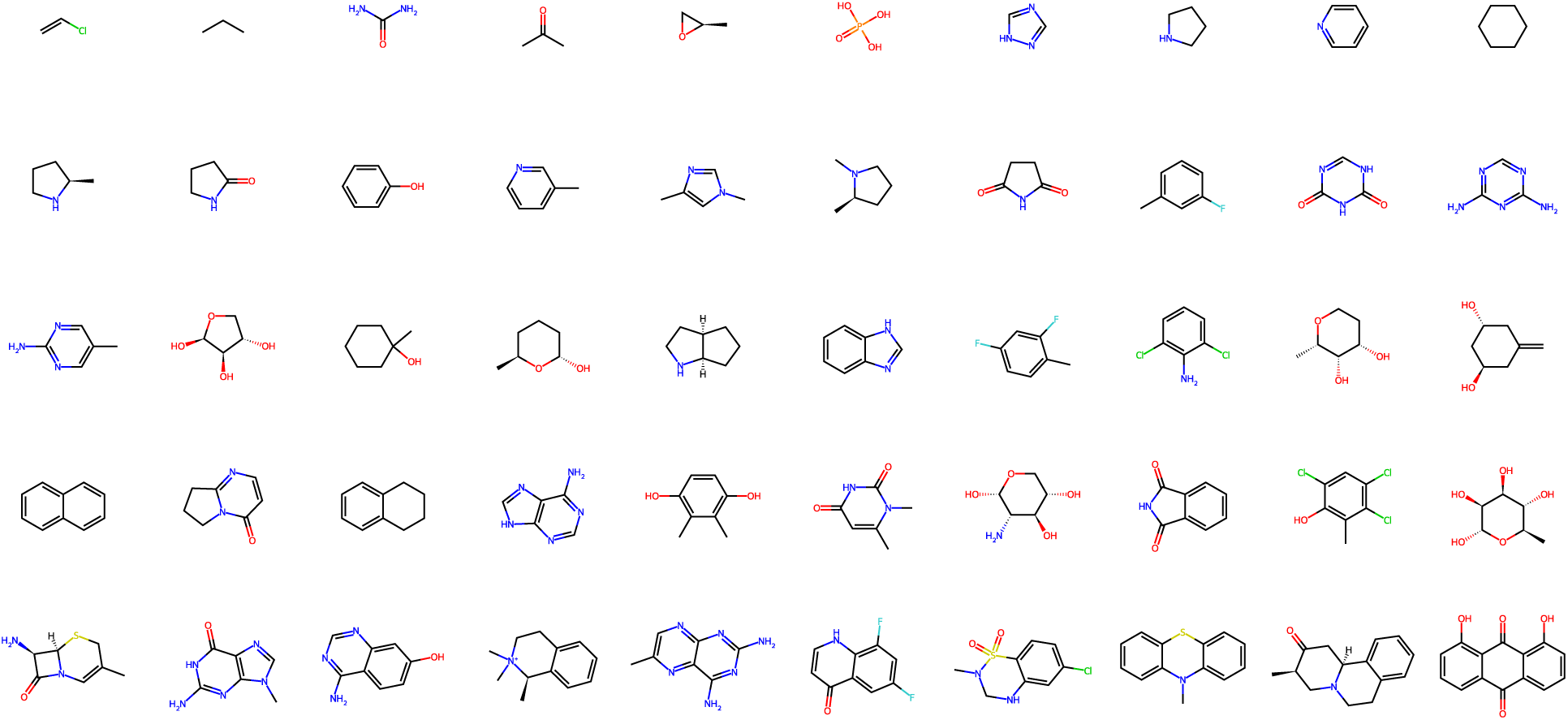
Structural formulas of the 50 representative fragments.

In COFFEE-PRESC, fragment pair queries of compound retrieval were generated based on 50 representative fragments, and we treat fragments with a similarity value of ≥ 0.45 as similar. Considering fragments derived from all actives across the 102 DUD-E targets, the fraction of fragment types that have a similar representative is 60.7%, while the fraction based on occurrences is 88.5%. These results indicate that, especially in terms of frequency, the representative set can cover most fragments that are likely to bind to target proteins.

### Accuracy of the docking simulation

Table 2 shows a summary of the mean enrichment factor (EF) values across the 102 DUD-E targets. Except for EF_1%_when pre-screening selects the top 2% of compounds, COFFEEPRESC achieves higher average EF values than Spresso in all cases.

**Table 2:**
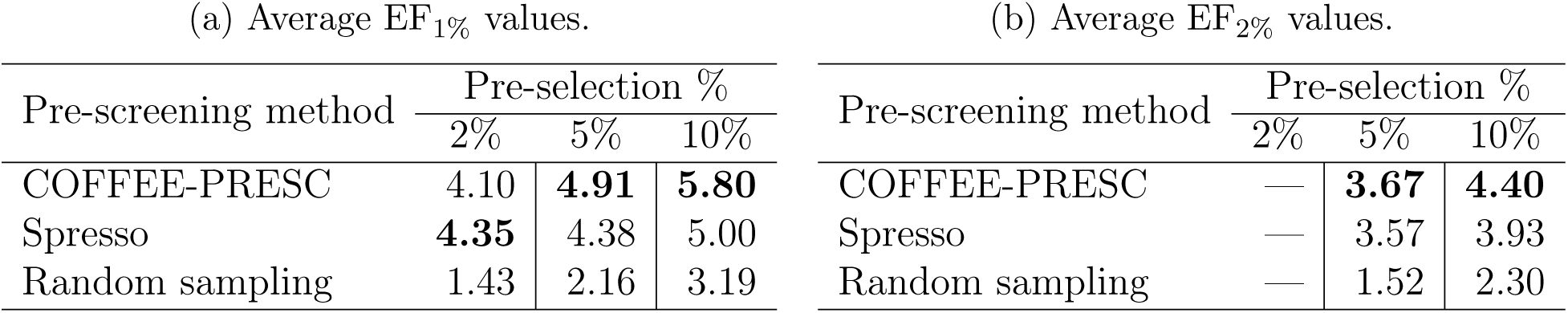
Average EF values for 102 DUD-E targets. The best value in each setting is shown in bold.

| (a) Average EF<sub>1%</sub> values. |  |  |  | (b) Average EF<sub>2%</sub> values. |  |  |  |
| --- | --- | --- | --- | --- | --- | --- | --- |
| Pre-screening method | Pre-selection % |  |  | Pre-screening method | Pre-selection % |  |  |
|  | 2% | 5% | 10% |  | 2% | 5% | 10% |
| COFFEE-PRESC | 4.10 | **4.91** | **5.80** | COFFEE-PRESC | — | **3.67** | **4.40** |
| Spresso | **4.35** | 4.38 | 5.00 | Spresso | — | 3.57 | 3.93 |
| Random sampling | 1.43 | 2.16 | 3.19 | Random sampling | — | 1.52 | 2.30 |

We examined the relationship between the prediction accuracy for each target and the coverage rate of fragment types constituting active compounds based on representative fragment sets. As shown in Figure S1, for all EF values across settings, the correlation coefficient was less than 0.1, indicating that the current number of representative fragments does not cause any meaningful decline in prediction performance. Given that COFFEE-PRESC achieves better prediction accuracy than Spresso, the current fragment set adequately represents important fragments required for accurate activity prediction.

### Virtual screening execution time

Table 3 shows the average CPU core time over 102 DUD-E targets. On average, each target contains approximately 14,000 compounds. COFFEE-PRESC, Spresso, and AutoDock Vina require approximately 0.21 CPU core hours, 6.69 CPU core hours, and 355.99 CPU core hours, respectively. In other words, COFFEE-PRESC is approximately 32 times faster than Spresso and approximately 1,685 times faster than AutoDock Vina.

**Table 3:** Average CPU core hours over 102 DUD-E targets.

| Screening method | Average execution time (CPU core h) | speedup relative to AutoDock Vina |
| --- | --- | --- |
| COFFEE-PRESC | 0.21 | $\times 1685$ |
| Spresso | 6.69 | $\times 53$ |
| AutoDock Vina | 355.99 | $\times 1$ |

Figure 5 shows plots of execution time versus the number of compounds, revealing that COFFEE-PRESC is consistently faster than Spresso and AutoDock Vina across all targets.

**Figure 5:**
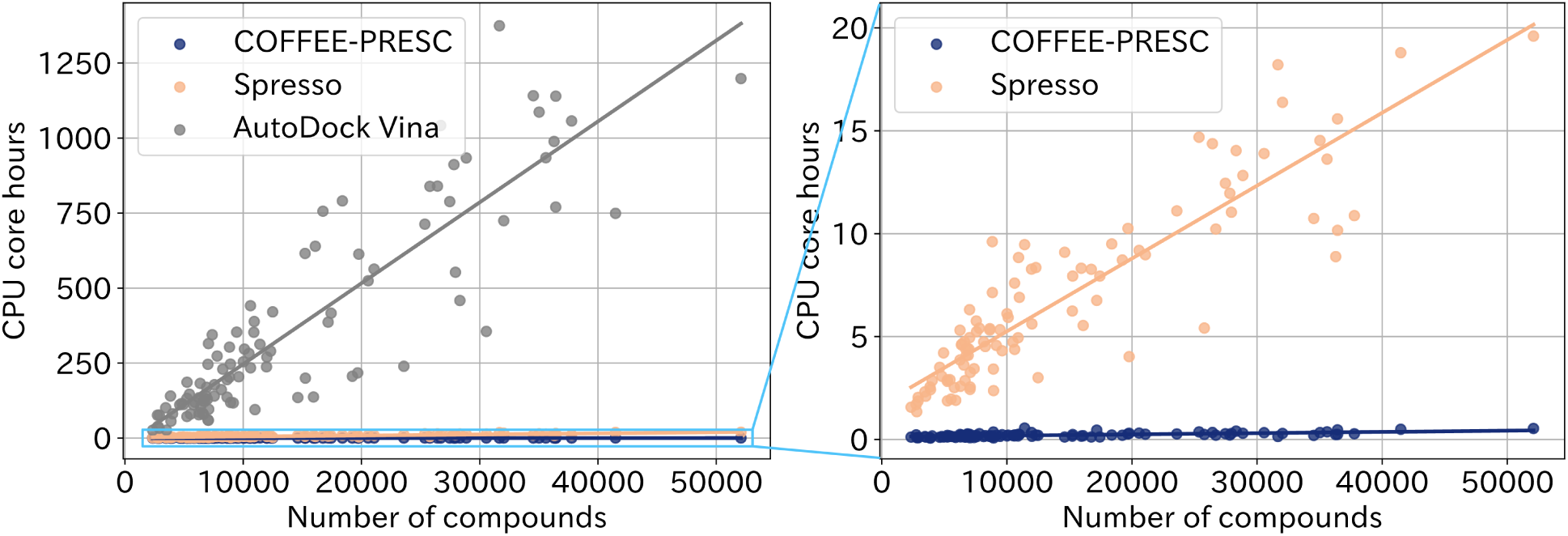
Relationship between the number of compounds and total execution time across 102 DUD-E targets. The left panel compares COFFEE-PRESC, Spresso, and AutoDock Vina, whereas the right panel focuses on COFFEE-PRESC and Spresso only. The straight line in each plot represents the regression line of the scatter plot.

## Discussion

### Coverage of fragments by representative fragment set

As described in the results section, the representative fragment set covered 88.5% of the active fragments in terms of occurrence, and 60.7% in terms of unique fragments. Figure 6 shows examples of fragments that are not covered (not similar to any representative fragment). The left and right fragments shown in Figure 6 were extracted from the ESR1 and KPCB targets, respectively. Some active compounds in the DUD-E dataset have fragments with four or more rings, including steroidal scaffolds such as the left fragment. In another case, macrocycles such as the right fragment are also not covered. Such large structures tend to exhibit high structural diversity because the number of derivative patterns increases with fragment size. Therefore, to ensure the representative fragment set covered a wider range of structures, large structures had to be excluded from the fragment population used for representative fragment selection. Since such large fragments are rare, we consider that they can be ignored, but the user may add such fragments to the representative fragment set according to prior knowledge, such as regarding the target protein.

**Figure 6:**
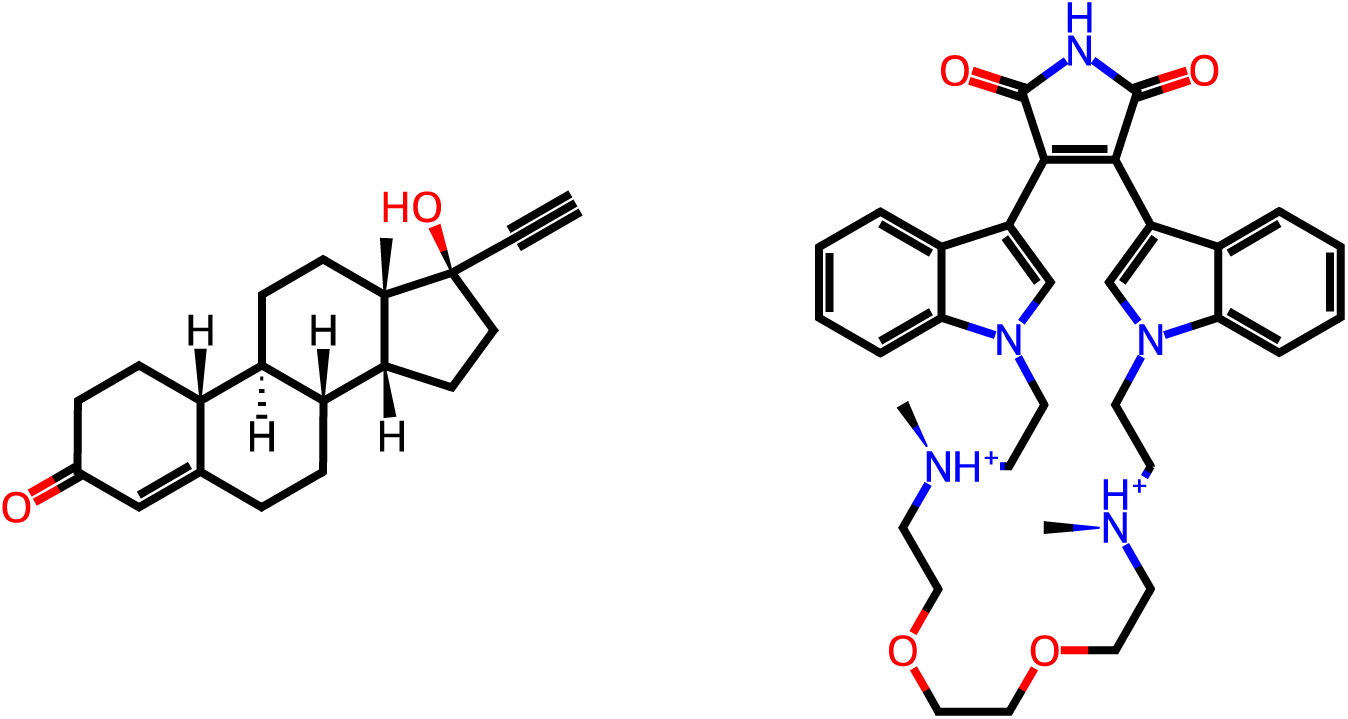
Examples of fragments from the DUD-E dataset that are not similar to any representative fragment. The left and right fragments were extracted from the ESR1 and KPCB targets, respectively.

### Effect of score penalty based on fragment similarity

COFFEE-PRESC performs similarity-based compound retrieval. Thus, matched compounds are scored not only by the docking scores of the fragment pair queries but also by penalties based on the similarity value in retrieval. To assess the effectiveness of the similarity-based penalty, we compared the prediction accuracy when using compound scores with and without the similarity-based penalty. Table 4 shows the average EFs over the 102 DUD-E targets in both settings. COFFEE-PRESC consistently achieves better performance when the fragment similarity value is considered for the penalty in the scoring function, indicating that the similarity-based penalty contributes to screening accuracy.

**Table 4:** Average EF values for 102 DUD-E targets with and without similarity-based penalty of compound scores in Eq. (2)

. The best value for each setting is shown in bold.
| (a) Average EF<sub>1%</sub> values. |  |  |  | (b) Average EF<sub>2%</sub> values. |  |  |  |
| --- | --- | --- | --- | --- | --- | --- | --- |
|  | Pre-selection % |  |  |  | Pre-selection % |  |  |
|  | 2% | 5% | 10% |  | 2% | 5% | 10% |
| Penalized score $E'$ | **4.10** | **4.94** | **5.80** | Penalized score $E'$ | — | **3.67** | **4.40** |
| Non-penalized score $E$ | 2.86 | 3.18 | 3.73 | Non-penalized score $E$ | — | 2.38 | 2.99 |

### Breakdown of execution time

To find prospects for further acceleration, we measured the execution time of each step in the COFFEE-PRESC workflow shown in Figure 1. Table 5 shows the average CPU time for Step 1 (pose exploration) and Step 3 (compound retrieval)―for example, three targets from the DUD-E dataset: PUR2, CP3A4, and EGFR2. These differ greatly in the number of compounds. Step 2 and Step 4 were not described here because their combined proportion of the total execution time is less than 5%.

**Table 5:** Breakdown of the execution time for each step of COFFEE-PRESC (CPU core seconds). Step 1 corresponds to docking pose exploration of representative fragments, and Step 3 corresponds to compound retrieval.

| Target | # compounds | Average execution time (CPU core s) |  |  |
| --- | --- | --- | --- | --- |
|  |  | Pose exploration<br>(Step 1) | Compound retrieval<br>(Step 3) | Total |
| PUR2 | 2,750 | 738 (91.8%) | 45 ( 5.6%) | 804 |
| CP3A4 | 11,970 | 978 (80.6%) | 207 (17.1%) | 1214 |
| EGFR | 35,592 | 752 (56.4%) | 555 (41.7%) | 1332 |

Step 1 corresponds to docking pose exploration of representative fragments. The same 50 representative fragments were used for all targets, leading to a computational complexity of *O*(|*F* |), where |*F* | is the number of representative fragments, and this cost does not depend on the number of ligand compounds |*L*|. Consequently, the per-compound execution time for Step 1 decreases as |*L*| increases.

Step 3 corresponds to compound retrieval. The number of compounds matching a retrieval is roughly proportional to the number of compounds in the database. Therefore, Step 3 scales with |*L*|, as retrieval and extraction processing are both affected by database size. As shown in Table 5, the contribution of Step 3 to total execution time grows with library size. Because Step 1 is independent of |*L*|, these results suggest that Step 3 will dominate execution time in ultra-large virtual screening. Since the computational cost in Step 3 is also influenced by the number of queries, further reducing them is the key to lowering the cost.

### Relationship between number of representative fragments and accuracy

We also verified how prediction accuracy changes with increasing the number of representative fragments. Table 6 shows the number of representative fragments (# *f^rep^*) and the corresponding COFFEE-PRESC EF values. Surprisingly, increasing the number of representative fragments from 50 to 100, 150 or 200 did not improve prediction accuracy in this experiment. We examined how the COFFEE-PRESC screening score changed from 50 to 200 representative fragments, The score value improved from an average of -5.84 to -6.67 for active compounds, and -5.28 to -6.25 for decoy compounds. Therefore, the decoy compounds, rather than active compounds, ended up improving the scores, causing the EF values to decrease. The higher the number of representative fragments, the more the likelihood of high similarity values. According to Eq. 2, the higher the similarity value, the better the score, which is why the score improves in this case. This suggests that structures exhibiting activity can be distinguished with only 50 fragments in this experiment. Also, adjusting the similarity threshold based on the number of representative fragments may improve accuracy.

**Table 6:** Average EF values for 102 DUD-E targets against COFFEE-PRESC compared by the number of representative fragments. The best EF value for each setting is shown in bold.

| (a) Average EF<sub>1%</sub> values. |  |  |  | (b) Average EF<sub>2%</sub> values. |  |  |  |
| --- | --- | --- | --- | --- | --- | --- | --- |
| # $f^{rep}$ | Pre-selection % | | | # $f^{rep}$ | Pre-selection % | | |
|  | 2% | 5% | 10% |  | 2% | 5% | 10% |
| 50 | **4.10** | **4.91** | **5.80** | 50 | — | **3.67** | **4.40** |
| 100 | 2.40 | 3.64 | 4.46 | 100 | — | 2.67 | 3.39 |
| 150 | 2.17 | 3.67 | 4.43 | 150 | — | 2.62 | 3.38 |
| 200 | 2.93 | 4.29 | 4.94 | 200 | — | 3.06 | 3.75 |

## Conclusion

In this study, we proposed COFFEE-PRESC, a pre-screening method that performs compound retrieval based on fragment docking pose pairs. By registering compounds in the library as sets of fragment pairs, COFFEE-PRESC enables fast compound evaluation while explicitly considering pairwise positional relationships between fragments. Furthermore, by using a small number of representative fragments that exhibit high similarity values to many others, COFFEE-PRESC reduces docking costs and performs similarity-based compound evaluation.

We evaluated the performance of COFFEE-PRESC on all 102 targets in the DUD-E dataset. Although the 50 representative fragments were selected regardless of their relationship with DUD-E, 88.5% of fragment occurrences in the dataset were covered. COFFEE-PRESC achieved higher accuracy than Spresso in 4 out of 5 EF settings. At the same time, on average, COFFEE-PRESC was 1685-fold faster than AutoDock Vina and 31-fold faster than Spresso. The relative advantage in execution time becomes more favorable with larger libraries, suggesting that COFFEE-PRESC can achieve faster pre-screening than Spresso for ultra-large virtual screening.

For further improvements in screening accuracy, it may be beneficial to retrieve compounds based on a greater number of fragments. This study focused on pairs of fragments to prioritize speed; however, relying on only two fragments may overlook important binding features. Extending the retrieval framework to fragment triplets or quadruplets could allow for more accurate compound evaluation when a compound contains many core fragments. However, increasing the number of fragments also increases the number of queries and database size, leading to a trade-off with computational cost. Addressing this trade-off through efficient query generation and indexing strategies will be an important direction for future work.

Overall, COFFEE-PRESC provides a promising foundation for accelerating SBVS by combining fragment-based representations with similarity-based retrieval. We anticipate that this approach will enable efficient exploration of the ever-growing chemical space in modern drug discovery.

## Data and software availability

Compound structures and protein structures were obtained from the Directory of Useful Decoys, Enhanced (DUD-E). We used OMEGA (version 5.0.0.3) to generate conformers and LigPrep (on Schrödinger suite, version 2024-1) to generate stereoisomeric and ionization states. The docking space of each target protein was determined using eBoxSize (version 1.1). The docking tool, AutoDock Vina (version 1.2.7), was used to evaluate the effectiveness of the proposed procedure. CSV files showing results for each DUD-E target and intermediate generated data for SAHH target have been deposited in Zenodo (https://zenodo.org/records/17644681, accessed on 8 December 2025).

## Acknowledgement

This study was carried out using the TSUBAME4.0 supercomputer at Institute of Science Tokyo. We extend our gratitude to OpenEye, Cadence Molecular Sciences for granting permission to use OMEGA for conformer generation. The authors thank Tomoya Saito, and Ayako Nunobe at Institute of Science Tokyo for the early stage studies of implementation of fragment search system, and fragment representative selection method, respectively.

## Supporting Information Available

The Supporting Information is available free of charge.

- Table S1: The 12 targets used to determine the parameters of COFFEE-PRESC.
- Figure S1: Relationship between coverage rate of fragment types constituing active compounds and EF values across 102 DUD-E targets.

## Author Information

### Authors

Masayoshi Shimizu, Department of Computer Science, School of Computing, Institute of Science Tokyo, Meguro-ku, Tokyo 152-8550, Japan; Satoshi Yoneyama, Department of Computer Science, School of Computing, Institute of Science Tokyo, Meguro-ku, Tokyo 152-8550, Japan; Keisuke Yanagisawa, Department of Computer Science, School of Computing, Institute of Science Tokyo, Meguro-ku, Tokyo 152-8550, Japan; orcid.org/0000-0003-0224-0035;

### Notes

The authors declare no competing financial interest.

## Abbreviations

AUC: Area under the curve
CP3A4: Cytochrome P450 3A4
CPU: Central processing unit
CSV: Comma-separated values
EF: Enrichment factor
EGFR: Epidermal growth factor receptor erbB1
ESR1: Estrogen receptor alpha
FBVS: Fragment-based virtual screening
FDA: Food and drug administration
KPCB: Protein kinase C beta
PUR2: GAR transformylase
RAM: Random access memory
RMSD: Root mean square deviation
SAHH: adenosylhomocysteinase
SBVS: Structure-based virtual screening
SMILES: Simplified molecular input line entry system.

