## Supplemental Table S1, Supplemental Figure S1 for "COFFEE-PRESC: A fast pre-screening method using compound retrieval by pairwise positional relationship of representative fragments"

Table S1: The 12 DUD-E targets used to determine the parameters of COFFEE-PRESC.

| Target | Average # fragments | |
| --- | --- | --- |
| | Actives | Decoys |
| ESR1 | 5.55 | 6.72 |
| GRIA2 | 6.47 | 6.52 |
| JAK2 | 6.23 | 6.63 |
| KIF11 | 6.78 | 6.18 |
| KPCB | 5.67 | 6.85 |
| PARP1 | 5.07 | 5.57 |
| PGH2 | 5.14 | 5.28 |
| PNPH | 3.93 | 6.08 |
| PRGR | 3.95 | 4.65 |
| PYRD | 5.66 | 5.64 |
| RXRA | 5.95 | 5.79 |
| SAHH | 3.46 | 5.57 |

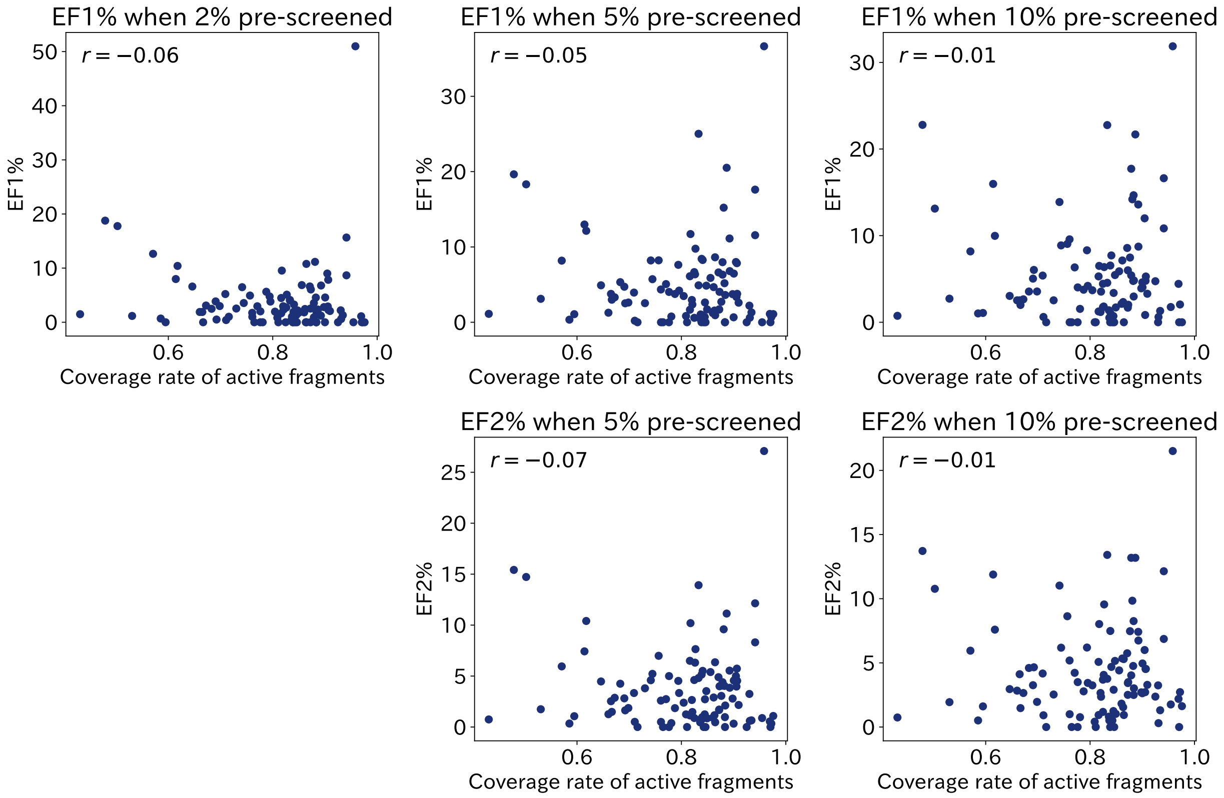


Figure S1: Relationship between coverage rate of fragment types constituing active compounds and EF values with COFFEE-PRESC across 102 DUD-E targets. The correlation coefficient is denoted as $r$.
